# Functional significance and therapeutic potential of miR-15a mimic in pancreatic ductal adenocarcinoma

**DOI:** 10.1101/716738

**Authors:** Shixiang Guo, Andrew Fesler, Wenjie Huang, Yunchao Wang, Jiali Yang, Xianxing Wang, Yao Zheng, Ga-Ram Hwang, Huaizhi Wang, Jingfang Ju

## Abstract

Treatment of pancreatic ductal adenocarcinoma (PDAC) remains a clinical challenge. There is an urgent need to develop novel strategies to enhance survival and improve patient prognosis. microRNAs (miRNAs) play critical roles as oncogenes or tumor suppressors in the regulation of cancer development and progression. In this study, we demonstrate that low expression of miR-15a is associated with poor prognosis of PDAC patients. miR-15a expression is reduced in PDAC while closely related miR-16 expression remains relatively unchanged. miR-15a suppresses several important targets such as Wee1, Chk1, Yap-1, and BMI-1 causing cell cycle arrest and inhibiting cell proliferation. Ectopic expression of miR-15a sensitizes PDAC cells to gemcitabine reducing the IC^50^ over 6.5-fold. To investigate the therapeutic potential of miR-15a, we used a modified miR-15a (5-FU-miR-15a) with uracil (U) residues in the guide strand replaced with 5-fluorouracil (5-FU). We demonstrated enhanced inhibition of PDAC cell proliferation by 5-FU-miR-15a compared to native miR-15a. *In vivo* we showed the therapeutic power of 5-FU-miR-15a alone or in combination with gemcitabine with near complete elimination of PDAC lung metastatic tumor growth. These results support the future development of 5-FU-miR-15a as a novel therapeutic agent as well as a prognostic biomarker in the clinical management of PDAC.

## Introduction

Pancreatic ductal adenocarcinoma (PDAC) is one of the most fatal malignancies leading to an estimated 45000 deaths each year, with an overall 5-year survival rate of less than 10% ^1^. Late clinical presentation, early metastasis and recurrence are mainly responsible for poor prognosis ^2^. Currently, the primary therapeutic approaches are curative resection, adjuvant chemotherapy and radiotherapy ^3-6^. Despite extensive research and development efforts aimed at enhancing the clinical management of PDAC, the outcomes remain unsatisfactory. For patients with metastatic disease five year survival remains below 5%, clearly demonstrating that current therapeutic strategies are not effective for improving patient outcomes ^1^. Hence, there is an urgent need to discover and develop novel diagnostic and therapeutic strategies for PDAC.

Evidence demonstrates that epigenetic regulation (methylation, non-coding RNAs) is a major mechanism contributing to cancer resistance. microRNAs (miRNAs) are a class of small non-coding RNAs (18-22 nucleotides) that play an important role in post-transcriptional regulation by binding to the 3’-UTR of target mRNA, leading to mRNA degradation or suppression of translation ^7-9^. The dysregulation of miRNAs is tightly correlated with proliferation, apoptosis, invasion, metastasis and resistance ^10, 11^. Recently, many studies have reported that miRNAs function as tumor suppressors or oncogenes in the pathogenesis and progression of PDAC ^12-14^.

miR-15a has been shown to be a major contributor in PDAC ^13, 15^. It was first reported that deletion of miR-15a/miR-16 is important in the development of chronic lymphocytic leukemia (CLL) ^16^. Recent studies have demonstrated that miR-15a acts as tumor suppressor by regulating a number of key targets (such as BMI-1, BCL2, Yap-1, DCLK1 etc.) in gastric cancer, colorectal cancer and PDAC ^12, 13, 17, 18^.

In this study, we demonstrated the clinical significance of miR-15a in PDAC as a prognostic biomarker based on our own patient cohort as well as validation based on The Cancer Genome Atlas (TCGA) database of PDAC. We identified and validated several important targets of miR-15a in PDAC. Ectopic expression of miR-15a causes cell cycle arrest and inhibits PDAC cell proliferation. miR-15a sensitizes PDAC cells to gemcitabine. We further demonstrated that a modified version of miR-15a (5-FU-miR-15a) has an enhanced impact on PDAC cell growth *in vitro* compared to native miR-15a. The 5-FU-miR-15a mimic was designed by replacing the uracil (U) residues of the guide stand of miR-15a with 5-fluorouracil (5-FU) ^17^. The rationale of such design is that we want to combine the therapeutic power of 5-FU and the tumor suppressive function of miR-15a into one entity ^17, 19^. 5-FU is a cornerstone chemotherapeutic agent for treating PDAC and many other solid tumors. This is a novel concept for design and development of miRNA based cancer therapeutics and we have demonstrated the therapeutic potential in colorectal cancer ^17, 19^. In this study, we show that 5-FU-miR-15a mimic is very potent, inhibiting PDAC metastatic tumor growth and sensitizing PDAC to gemcitabine *in vivo*. As a result, 5-FU-miR-15a has great potential to be further developed as a novel therapeutic drug for PDAC.

## Results

### The expression of miR-15a is significantly associated with PDAC patient survival

To determine the expression level of miR-15a in PDAC patients and its relationship with patient survival, levels of miR-15a and miR-16 were quantified by qRT-PCR analysis using normal (10) and PDAC (30) tissue samples. Our results showed that miR-15a expression was significantly (P=0.0014) decreased in PDAC tissue compared to normal pancreas, while there was no considerable alteration of miR-16 expression between PDAC tissue and normal pancreas (P=0.4118) (Figure 1A). In addition, Kaplan-Meier survival analysis of miR-15a expression and PDAC patient survival based on TCGA PDAC RNA-seq expression database shows that the overall survival time of patients with low-expression of miR-15a was significantly shorter (p<0.0001) compared to patients with high-expression of miR-15a (Figure 1B). The median survival time is 511 days and 1502 days respectively. It has been demonstrated that miR-15a and miR-16 are clustered on Chr13q14 (Figure 1C) ^16^. To determine the potential mechanism of reduction of miR-15a expression in PDAC, we used primers to distinguish pri-miR-15a/16, pre-miR-15a vs. pre-miR-16, and mature miR-15a vs. mature miR-16 and measured their expression in both the nucleus and the cytoplasm (Figure S1A). Our results show that there is no difference between pre-miR-15a and pre-miR-16. However, expression of mature miR-15a in PDAC was significantly lower than expression of miR-16 (Figure 1D). In addition, we tested the expression levels of primary, precursor and mature miR-15a and miR-16 in clinical specimens (16 PDAC tissues and 8 normal pancreas tissues) using real-time PCR. The results showed that mature miR-15a was significantly reduced in expression compared to miR-16, while there were no differences between precursor miR-15a and precursor miR-16 in clinical samples (Figure 1E). The results are consistent with those of three PDAC cell lines (Figures 1D & S1B). These results suggest that the reduction of miR-15a in PDAC occurs during the processing of pre-miR-15a to mature miR-15a.

**Figure 1.**
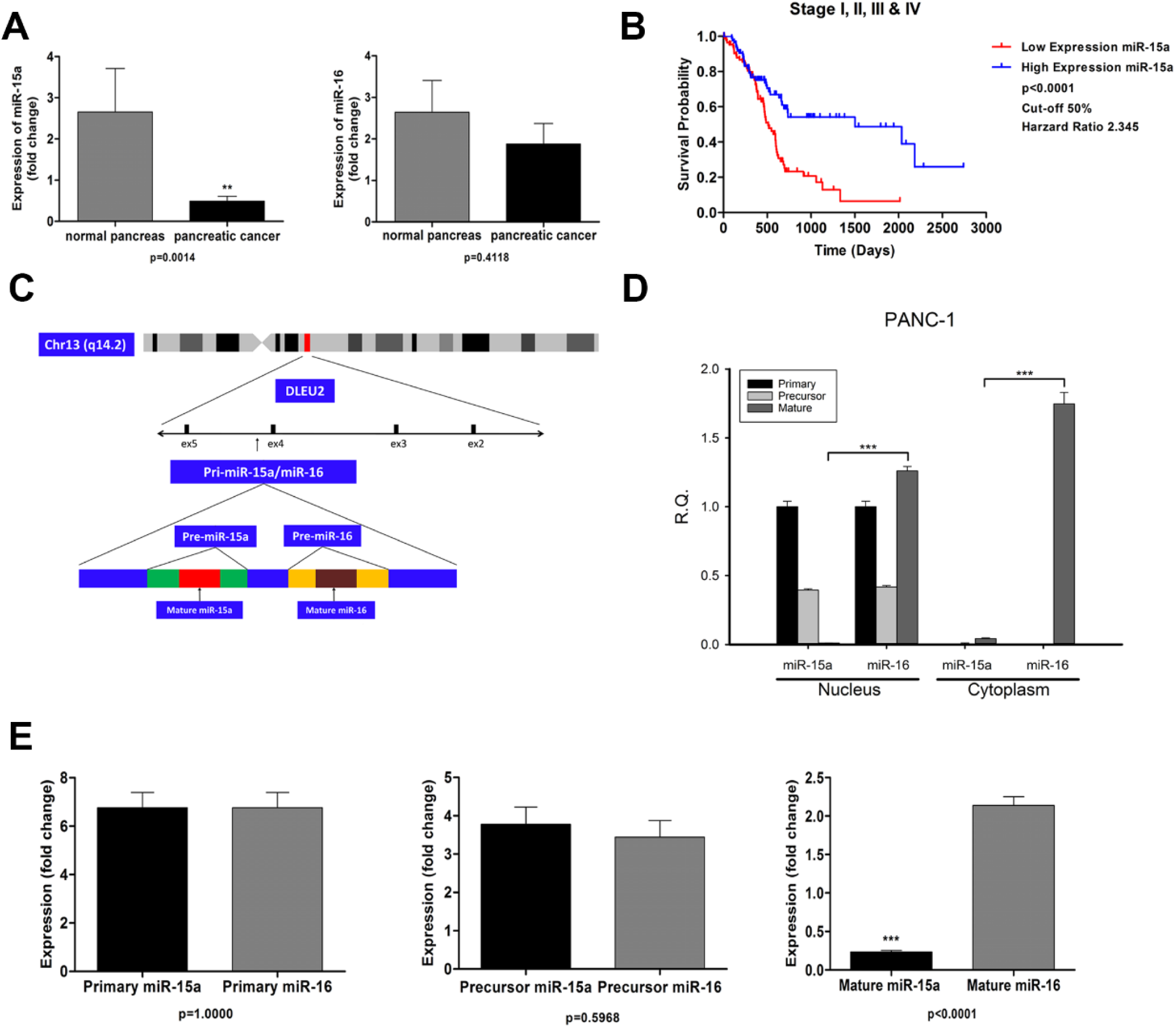
miR-15a expression is reduced in PDAC while miR-16 expression remains unchanged. **(A)** miR-15a expression is significantly decreased in PDAC compared to normal pancreas (p=0.0014), while there is no considerable difference in miR-16 expression between PDAC tissue and normal pancreas. (p=0.4118) **(B)** Low expression of miR-15a is associated with worse prognosis for PDAC patients. (*** p<0.001) (HR: 2.345) **(C)** miR-15a and miR-16 are both located on Chr13q14.2 They are transcribed as one primary transcript that is then processed into pre-miR-15a and pre-miR-16. These precursors are then further processed into mature miR-15a and miR-16 **(D)** In PANC-1 cells expression of pre-miR-15a and pre-miR-16 are similar in both the nucleus and the cytoplasm. Expression of mature miR-16 is significantly higher than miR-15a in both the nucleus and the cytoplasm. (N=3) **(E)** Mature miR-15a expression is significantly reduced compared to miR-16 (p<0.0001), while there are no difference between primary and precursor of miR-15a/miR-16 in PDAC tissues.

### Inhibition of PDAC cell proliferation, cell cycle arrest, and chemosensitization to gemcitabine by ectopic expression of miR-15a

To investigate the impact of miR-15a on pancreatic cancer cell proliferation, we transfected native miR-15a, 5-FU-miR-15a or control miRNA at 50 nM into three pancreatic cancer cell lines (AsPC-1, PANC-1 and Hs 766T), and a WST-1 proliferation assay was performed to measure cell viability. Our results show that miR-15a inhibits the proliferation of pancreatic cancer cells (Figure 2A). The inhibitory effect of miR-15a was enhanced by modified 5-FU-miR-15a mimic in AsPC-1 (IC^50^=14.7255 nM), PANC-1 (IC^50^=23.4587 nM) and Hs 766T (IC^50^=18.713 nM) (Figure 2A). We further demonstrated 5-FU-miR-15a has the therapeutic power of both miR-15a and 5-FU using 5-FU modified *C. elegans* cel-miR-67 as control (Figure 2B). 5-FU-cel-miR-67 treated cells only exhibit the effect of 5-FU alone, while the therapeutic effect of miR-15a is missing.

**Figure 2.**
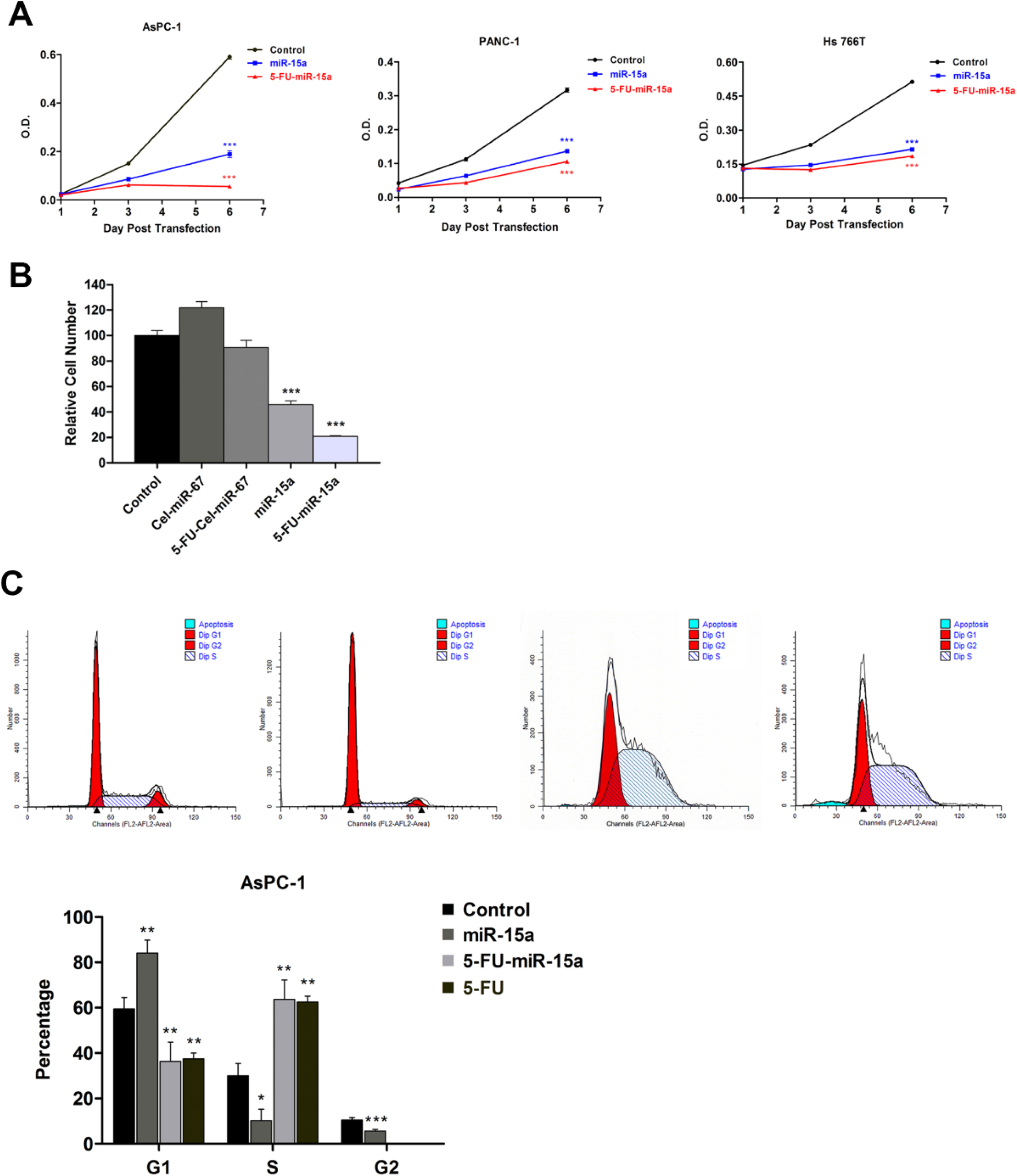
miR-15a and 5-FU-miR-15a inhibit PDAC cell proliferation. **(A)** 5-FU-miR-15a is more effective at suppressing pancreatic cancer cell proliferation compared native miR-15a in three pancreatic cancer cell lines. (* p<0.05, ** p<0.01, *** p<0.001)(N=3) **(B)** Cell number 6 days after transfection was decreased 10% by 5-FU modified control miRNA, *C. elegans* cel-miR-67 (5-FU-Cel-miR-67), while 5-FU-miR-15a reduced cell number by 80%. (*** p<0.001)(N=3) **(C)** 5-FU-miR-15a induces the alteration of cell cycle progression (increasing S phase and G2 disappearance). Unmodified miR-15a induces G1 arrest and 5-FU alone induced S phase arrest similar to 5-FU-miR-15a. (* p<0.05, ** p<0.01, *** p<0.001)(N=3)

We analyzed the impact of both miR-15a and 5-FU-miR-15a on cell cycle by flow cytometry. Our results show that miR-15a triggered cell cycle arrest at G1 phase with a significant reduction in G2 phase (Figure 2C). The 5-FU alone treated PDAC cells exhibited increased S phase as a result of DNA damage response. We also observed the increased S phase in 5-FU-miR-15a mimic treated cells, in addition to G2 check point reduction. Western immunoblotting showed that there is a reduction of cyclin A expression along with increased levels of p27 in cells treated with miR-15a (Figure S2).

We also found that 5-FU-miR-15a sensitizes PDAC cells to gemcitabine. 5-FU-miR-15a has a synergistic effect when combined with gemcitabine to enhance the therapeutic effect in pancreatic cancer cells (Figure 3A). The synergistic effect was analyzed using CombuSyn software to reveal that there was a synergistic effect of 5-FU-miR-15a combined with gemcitabine at lower concentrations (CI value < 0.4). In combination, the IC^50^ of 5-FU-miR-15a was reduced by over 3-fold from 18.713 to 5.886 nM, and the IC^50^ of gemcitabine was decreased by over 6-fold from 3.876 to 0.588 μM (Figure 3B). Overall, 5-FU-miR-15a enhanced the therapeutic effect of gemcitabine on pancreatic cancer cells.

**Figure 3.**
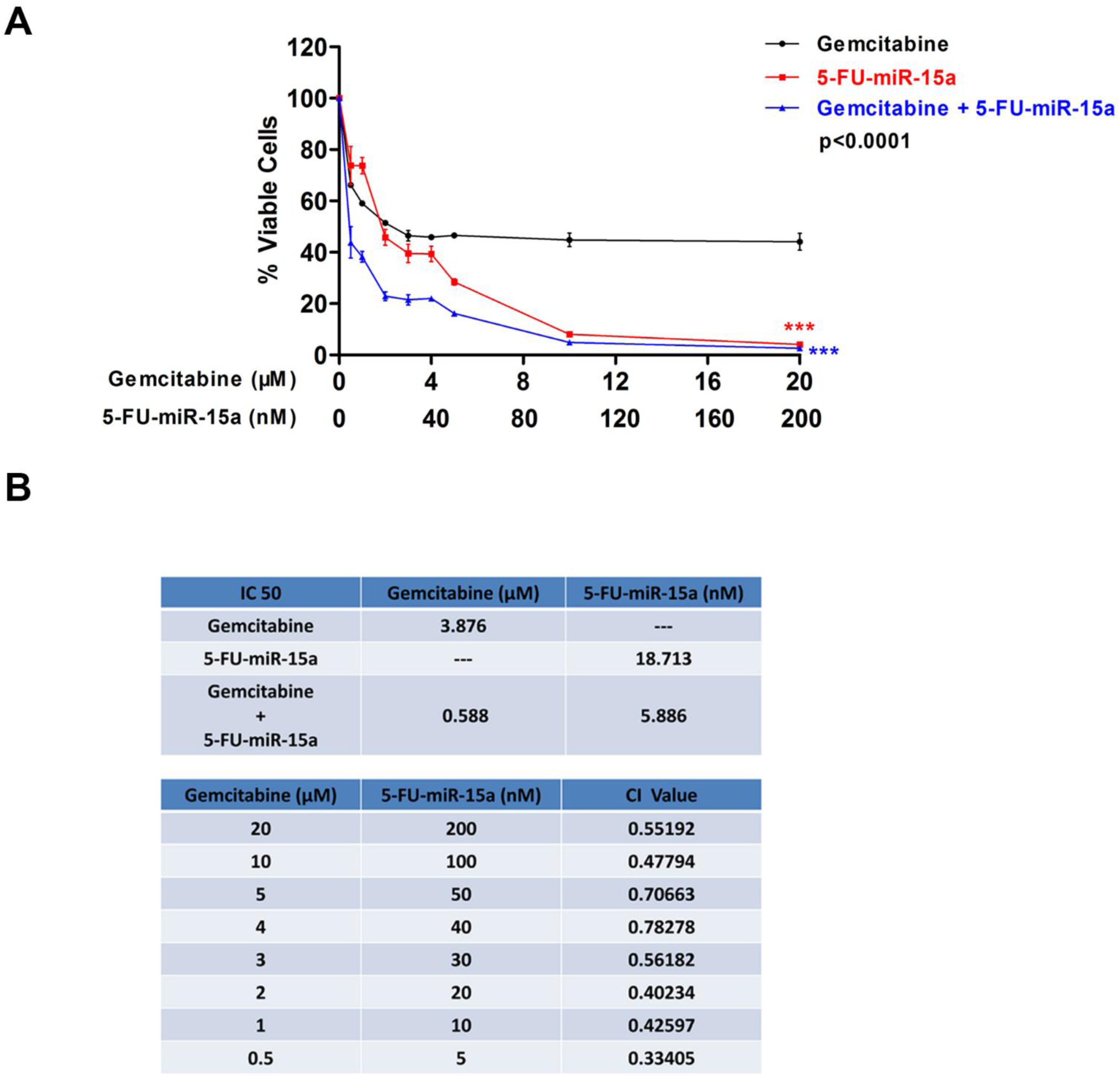
5-FU-miR-15a increases PDAC sensitivity to gemcitabine. **(A)** Gemcitabine and 5-FU-miR-15a inhibit PDAC cell viability and the effects are enhanced when gemcitabine is combined with 5-FU-miR-15a. (*** p<0.001) **(B)** CI values calculated for 5-FU-miR-15a and gemcitabine demonstrate that there is a significant synergistic effect at low concentrations. In combination, the IC^50^ of 5-FU-miR-15a is reduced from 18.713 to 5.886 nM, and the IC^50^ of gemcitabine is reduced from 3.876 to 0.588 μM.

### miR-15a suppressed the expression of several key targets Wee1, Chk1, BMI-1, and Yap-1 in PDAC

To understand the molecular mechanism and functional impact of miR-15a in PDAC, we investigated mRNA targets of miR-15a. Using TargetScan and miRDB, we discovered several potential targets of miR-15a, including Wee1 and Chk1, both genes are important in regulating G2 cell cycle control ^20^. Wee1 and Chk1 both have two potential miR-15a binding sites in their 3’UTR’s (Figure 4A).

**Figure 4.**
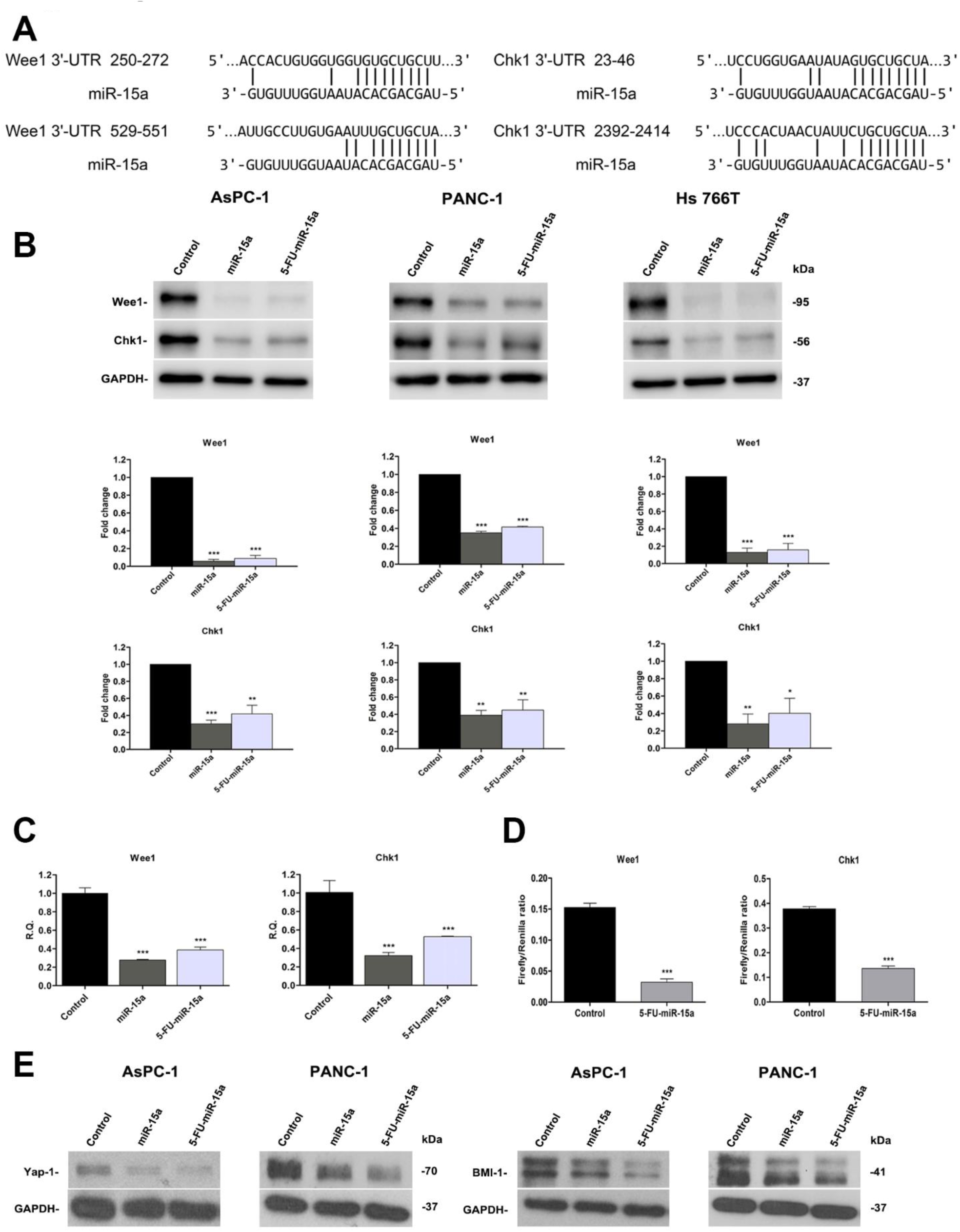
miR-15a and 5-FU-miR-15a inhibit the expression of several important targets in PDAC. **(A)** There are two potential binding sites for miR-15a in the 3’UTR’s of both Wee1 and Chk1 **(B)** Western blot analysis demonstrates miR-15a and 5-FU-miR-15a significantly inhibit expression of target genes Wee1 and Chk1. (* p<0.05, ** p<0.01, *** p<0.001)(N=3) **(C)** qRT-PCR analysis shows that miR-15 and 5-FU-miR-15a inhibit Wee1 and Chk1 expression at the mRNA level. (*** p<0.001)(N=3) **(C)** The luciferase assay confirms 5-FU-miR-15a and miR-15a directly bind to the 3’UTR’s of Wee1 and Chk1. (*** p<0.001)(N=3) **(D)** miR-15a and 5-FU miR-15a inhibit expression of previously identified targets Yap-1 and BMI-1(N=3)

Western immunoblotting was used to quantify the expression of Wee1 and Chk1 after ectopic expression of miR-15a in PDAC cell lines. Our results show there is a significant reduction of Wee1 and Chk1 expression with restoration of miR-15a and 5-FU-miR-15a. The quantification of Wee1 and Chk1 expression demonstrated that the reduction was significant and ranged from 70% to 95% (Wee1, miR-15a), 60% to 90% (Wee1, 5-FU-miR-15a), 65% to 75% (Chk1, miR-15a), 55% to 60% (Chk1, 5-FU-miR-15a) respectively (Figure 4B). We used Wee1 and Chk1 specific siRNAs as positive controls for reduction of Wee1 and Chk1 expression (Figure S3A). These results also confirmed that 5-FU-miR-15a mimic retained target specificity for Wee1 and Chk1. Transfection with 5-FU-cel-miR-67 had no effect on expression of Wee1 or Chk1 (Figure S3B). In addition, we also quantified the mRNA levels of Wee1 and Chk1 after transfection of miR-15a into PDAC cell lines by real-time PCR. The results showed that the mRNA levels of these two targets were both decreased in pancreatic cancer cells transfected with miR-15a and 5-FU-miR-15a (Figure 4C). In addition, to further validate miR-15a directly binding to the 3’-UTR of Wee1 and Chk1 mRNA, we performed a luciferase reporter assay. As shown in Figure 4D, miR-15a and 5-FU-miR-15a can both significantly reduce luciferase activity compared to control.

We also confirmed that two previously identified targets of miR-15a in colon cancer, BMI-1 and Yap-1 ^17^, were reduced by miR-15a in PDAC. The 5-FU modification of 5-FU-miR-15a did not alter target specificity for both BMI-1 and Yap-1 (Figure 4E).

### Cdc2 is the downstream target of Wee1 and correlated with PDAC patient survival

Cdc2 is an important downstream target of the Wee1 signaling pathway in PDAC ^21^. Therefore, we used the clinical data from TCGA database to analyze the relationship between Cdc2 expression and patient prognosis. We downloaded clinical data for 176 PDAC patients’ and performed survival analysis. The result show the overall survival time of patients with high Cdc2 expression was shorter than patients with low Cdc2 expression (Figure 5A). The median survival time is 702 days and 498 days respectively. In addition, we extracted 101 PDAC patients’ FFPE tissues and 10 normal pancreas FFPE tissues from our clinical sample center, and performed IHC to detect Cdc2 expression. IHC analysis showed that Cdc2 was over-expressed in PDAC tissue compared to normal pancreas (Figure 5B).

**Figure 5.**
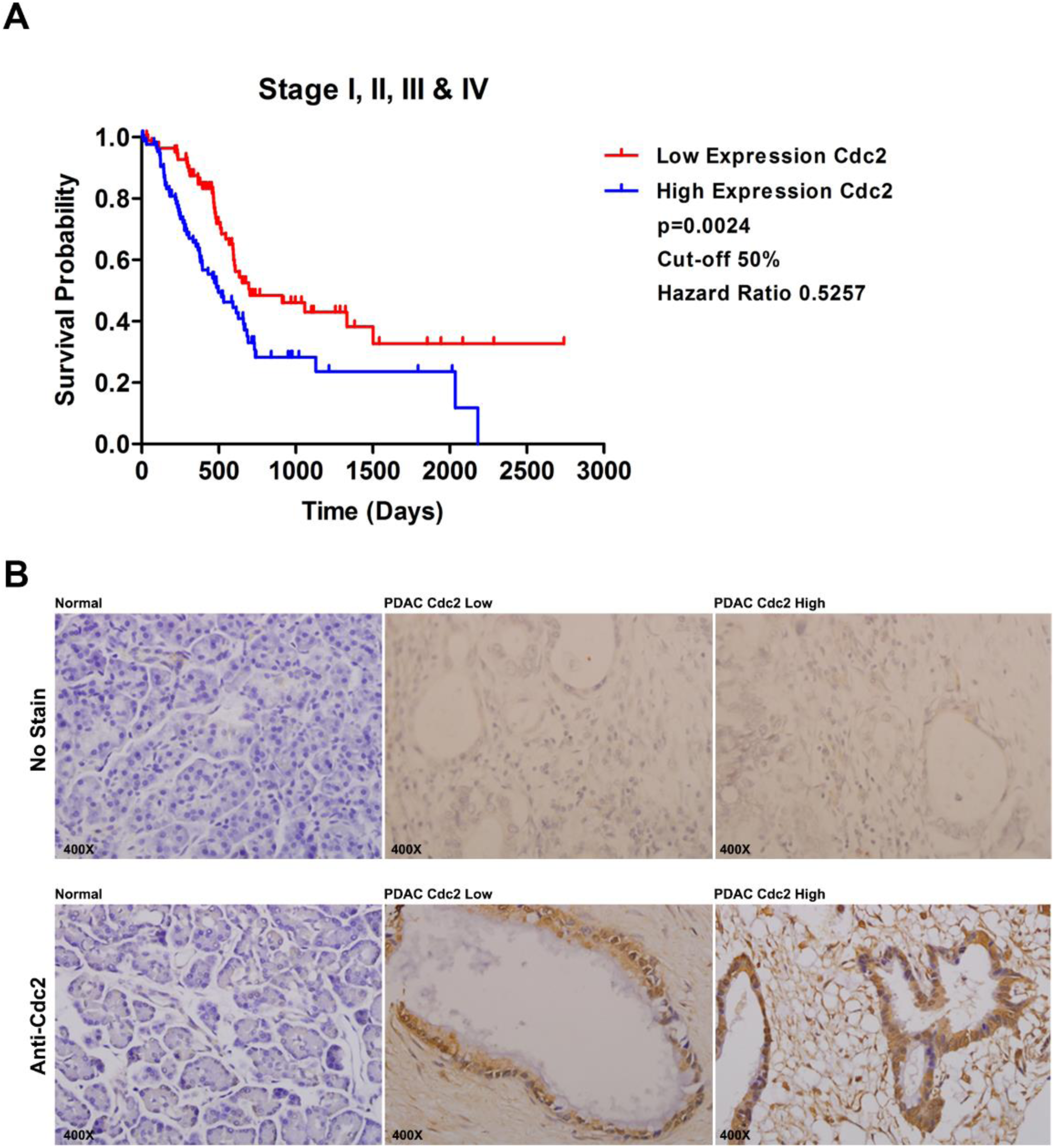
Wee1 target Cdc2 is a negative prognostic marker for PDAC. **(A)** High expression of Cdc2 is associated with worse prognosis for PDAC patients. (p=0.0024)(HR: 0.5257) **(B)** Cdc2 is not expressed in normal pancreas tissue and Cdc2 is over-expressed in pancreatic cancer tissues.

### 5-FU-miR-15a suppresses metastatic tumor formation and overcomes chemoresistance

Patients with PDAC succumb to the disease due to tumor metastasis. To investigate the impact of 5-FU-miR-15a on pancreatic cancer metastasis in *vivo*, we established a pancreatic cancer mouse metastasis model via tail vein injection of metastatic pancreatic cancer cells (Hs 766T). Three days after establishing metastatic tumors, we divided mice into four groups and started treatment. The results showed that 5-FU-miR-15a (3.2 mg/kg) significantly inhibited metastatic tumor growth at a dose that was 15-fold less than that of gemcitabine (50mg/kg) and the inhibitory effect was enhanced in combination with gemcitabine (Figure 6A &B). The treatment was stopped on day 17 and resumed on day 32 to determine whether mice treated with 5-FU-miR-15a develop resistance. While mice treated with gemcitabine developed resistance, we did not observe any significant resistance in mice after we resumed treatment with 5-FU-miR-15a for 4 additional injections (P=0.0179) (Figure 6C). All mice treated with 5-FU-miR-15a or gemcitabine displayed no signs of toxicity. Furthermore, we tracked survival time of all mice and performed survival analysis. We found that survival time of mice treated with 5-FU-miR-15a was significantly (P = 0.0257) longer than control mice (76 days vs. 52 days). In combination with gemcitabine the effect was more pronounced (92 days vs. 52 days) (P=0.0021) (Figure 5D).

**Figure 6.**
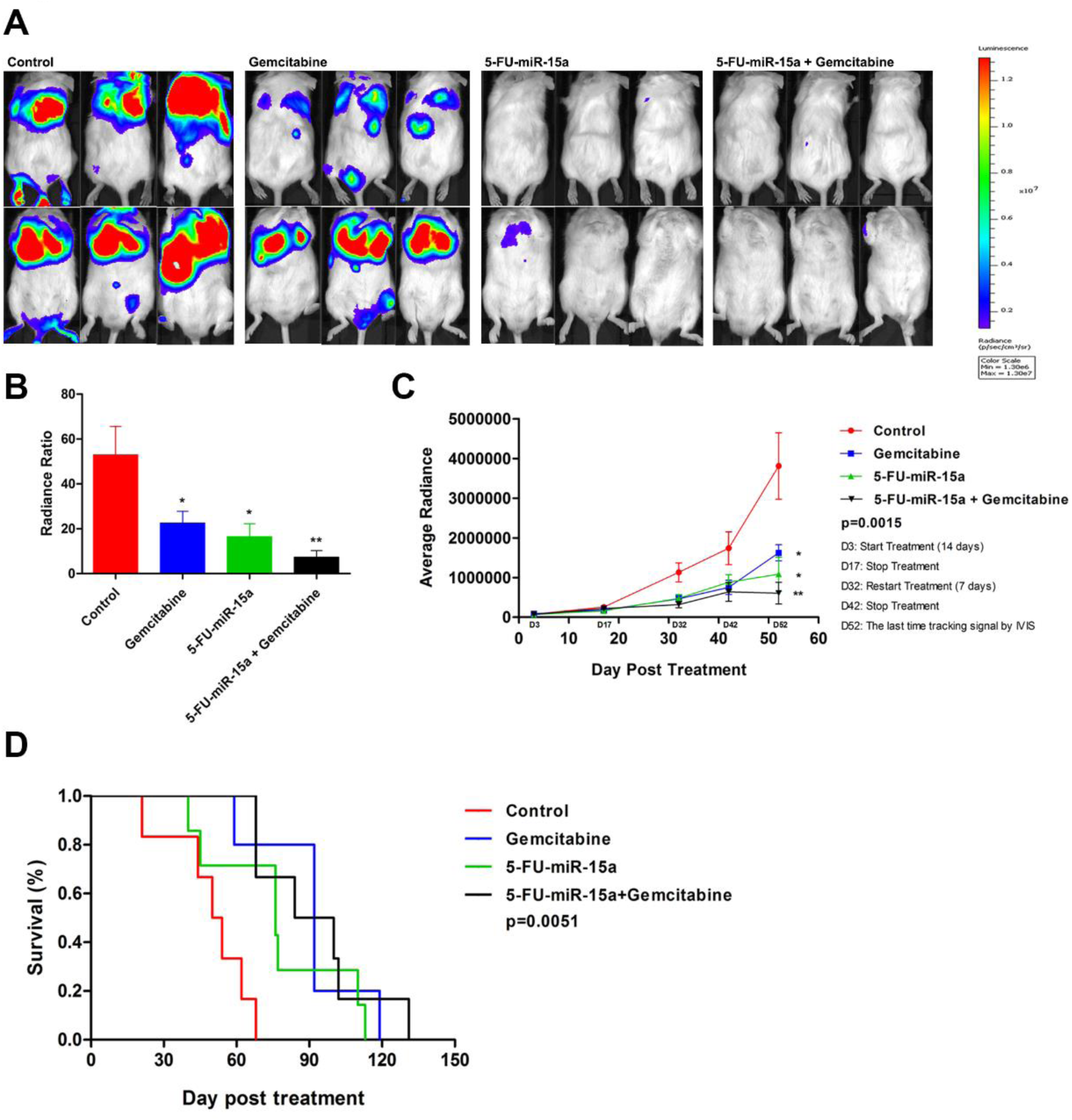
5-FU-miR-15a inhibits PDAC metastatic tumor growth *in vivo*. **(A)** Representative images show 5-FU-miR-15a (3.2 mg/kg) inhibits metastatic tumor growth at a dose that is 15-fold less than that of gemcitabine (50 mg/kg) and the inhibitory effect is enhanced in combination with gemcitabine. **(B)** Quantification of the measured radiance from the end of the experiment (Day 52) vs the start (Day 3) shows that effective inhibition of tumor growth by gemcitabine, 5-FU-miR-15a and 5-FU-miR-15a + gemcitabine. (* p<0.05)(N=6) **(C)** Quantified IVIS measurements through the course of the experiment show the inhibitory effects of 5-FU-miR-15a. 5-FU-miR-15a significantly inhibits metastatic PDAC tumor growth. There is no resistance in mice treated with 5-FU-miR-15a, while mice treated with gemcitabine developed resistance. (* p<0.05, ** p<0.01)(N=6) **(D)** 5-FU-miR-15a extends survival time compared to mice treated with control miRNA (76 day vs. 52 day median survival) (P=0.0257) (HR: 2.851). Combination of 5-FU-miR-15a increased median survival to 92 days (P=0.0021) (HR: 4.071)(N=6).

## Discussion

In this study, we discovered that reduced expression of miR-15a in PDAC was significantly associated with poor patient survival based on TCGA database. The results suggest miR-15a has potential as a prognostic biomarker in PDAC. This is highly consistent with our previous studies that miR-15a was a prognostic biomarker in PDAC ^13^. One novel aspect of the study is the experimental evidence that reduction of mature miR-15a in PDAC occurs during processing of pre-miR-15a to mature miR-15a as closely related miR-16 expression level is not altered in PDAC. We also discovered a similar expression pattern of reduced miR-15a in gastric and colorectal cancer, while miR-16 can be used as a housekeeping miRNA ^18, 22^. This is a unique mechanism for loss of miR-15a expression compared to miR-15a deletion in CLL ^23^. Based on the miRNA biogenesis pathway, pre-miRNA is cleaved to generate a mature duplex miRNA by Dicer after pre-miRNA is exported into the cytoplasm by Exportin-5 ^11, 24^. Dicer cooperates with other proteins to influence miRNA maturation, and its dysregulation leads to the alteration of mature miRNA expression ^25^. However, the exact nature of such differential processing of miR-15a and miR-16 in PDAC remains to be further investigated in future studies.

miR-15a was initially discovered in CLL as a tumor suppressor ^16^. miR-15a has also been extensively investigated in various tumor types as a tumor suppressor, targeting a number of important genes ^13, 17, 26, 27^. In this study we demonstrated miR-15a functions as a tumor suppressor in PDAC *in vitro* by inhibiting cell proliferation and impacts cell cycle control. We discovered and validated several important targets of miR-15a in PDAC such as Wee1, Chk1, BMI-1 and Yap-1. It has been well documented that the expression of all of these targets is elevated in PDAC and many are good target candidates for therapeutic development in PDAC. BMI-1 is an oncogene and over expression is associated with poor prognosis ^18, 28^. BMI-1 promotes PDAC invasion and metastasis and inhibition of BMI-1 enhances gemcitabine sensitivity ^29, 30^. Yap-1 is crucial in promoting pancreatic tumorigenesis ^31^. Yap-1 is activated in p53 deficient tumors and p53 is mutated in about 75% of PDAC patients ^32^. Yap-1 expression promotes cell cycle proliferation as well as invasion and migration. Yap-1 expression is also associated with chemoresistance and poor prognosis ^33, 34^. Wee1 and Chk1 are two key G2/M checkpoint regulators, which can affect Cdc2 activity ^35, 36^. These targets have been recognized as candidates for therapeutic development by the pharmaceutical industry ^20^. However, the results have been disappointing as single targeted approaches such as LY2603618 which suppresses the expression of Chk1, failed to yield any synergy with gemcitabine in a phase II clinical trial for PDAC ^37^. While it has been demonstrated that the combination of gemcitabine with Wee1 inhibitor, MK-1775, results in enhanced tumor regression compared to gemcitabine alone in p53-deficient tumors ^38^. MK-1775 itself failed to trigger any tumor regression in PDAC. It has been shown recently by Chuang *et al*. that there is potential for targeting cell cycle checkpoint functions such Wee1 and Chk1 in PDAC ^35^. The drug screening effort in this study clearly showed that the combination of gemcitabine with Chk1 and Wee1 inhibition provided strong disease control in all tumor xenograft models ^35^. These findings along with the other targets we validated, demonstrate the unique potential of developing miR-15a mimic as a therapeutic agent as it can suppress a number of important targets to impact multiple pathways (Figure 7). In addition to the targets we have assessed in this study, previously identified targets of miR-15a such as BCL2 and DCLK1 have important functions in PDAC ^17, 23^. BLC2 expression is associated resistance to apoptosis and metastatic potential of PDAC and may be a potential therapeutic target ^39, 40^. DCLK1 is a marker for stem like pancreatic cancer cells and promotes EMT ^41, 42^. DCLK1 expression is also associated with tumor progression, worse patient survival and DCLK1 inhibition increases the effectiveness of gemcitabine ^43-45^. The multi-targeted function of miR-15a is a major advantage for cancer drug development as it offers a unique mechanism to overcome resistance associated with many of the previously developed targeted therapies. This strategy creates a new drug that is better than the simple combination of 5-FU with miR-15a. The combination approach will be hampered by 5-FU’s toxic side effects. Through incorporation of 5-FU into miR-15a we achieve potency at a much lower concentration avoiding toxicity. We initially implemented this strategy using miR-129 and miR-15a in colorectal cancer ^17, 19^. In this study, we found that 5-FU-miR-15a maintains target specificity and is more effective than unmodified miR-15a.

**Figure 7.**
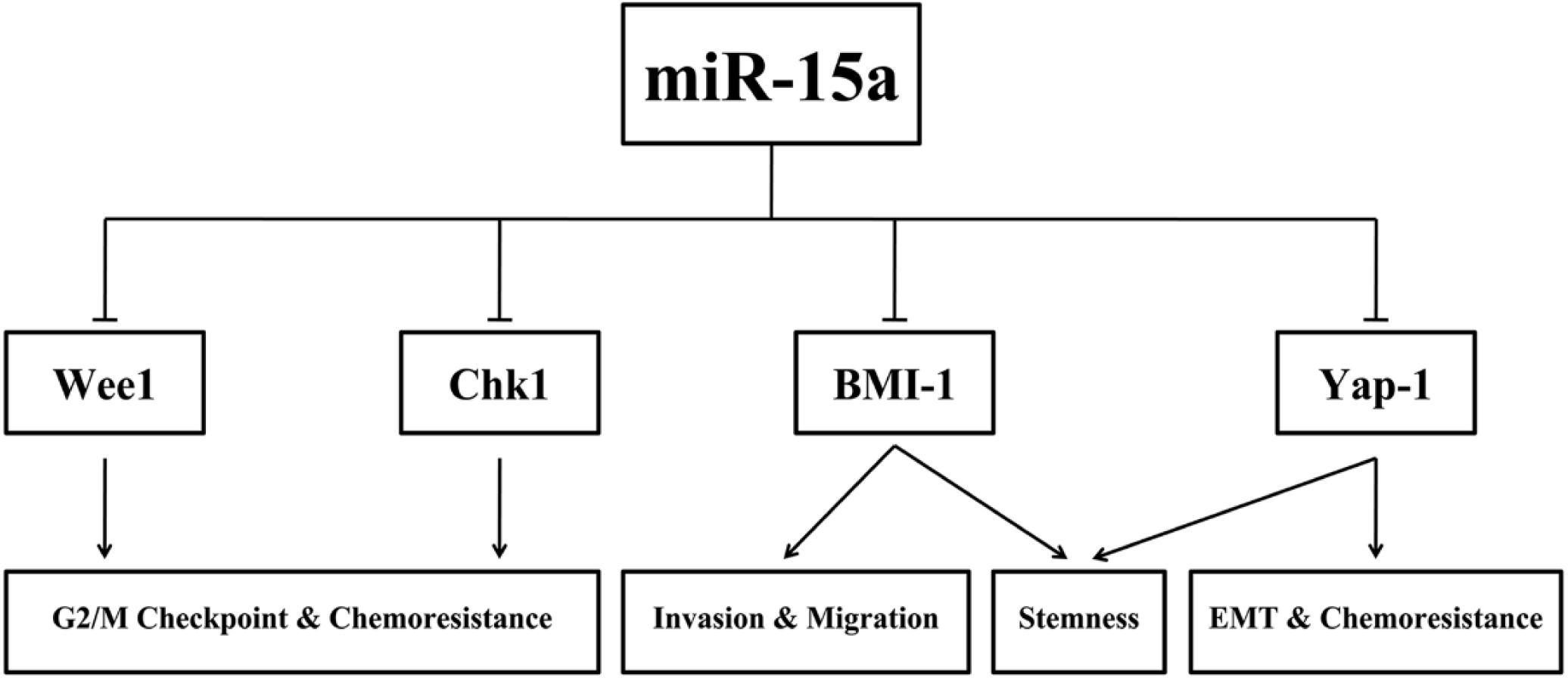
Schematic of important miR-15a targets in PDAC and the pathways they regulate. miR-15a inhibits several important targets in PDAC including Wee1, Chk1, BM1-1 and Yap-1. Through its inhibition of these targets, miR-15a is able to induce cell cycle arrest, enhance chemosensitivity, and inhibit invasion and migration.

Finally, we determined the therapeutic impact of 5-FU-miR-15a mimic *in vivo* using a tail vein injection PDAC metastasis model. As PDAC metastasis in lung and liver is the major cause of patient death, this mouse model is simple and effective in evaluating the therapeutic impact of drugs for PDAC ^46, 47^. To demonstrate the proof-of-concept, in a pancreatic cancer mouse metastasis model, we showed that 5-FU-miR-15a (at 3.2 mg/kg) markedly inhibited metastatic tumor growth at a dose that is 15-fold less than that of gemcitabine (50 mg/kg), and the inhibitory effect of lung tumor growth was further enhanced in combination with gemcitabine, consistent with the synergistic effect we observed *in vitro*. While mice treated with gemcitabine alone develop resistance, we did not observe any resistance in mice treated with 5-FU-miR-15a. Importantly, all mice treated with 5-FU-miR-15a displayed no signs of systemic toxicity. The lack of toxicity will be a major advantage for developing effective anticancer therapy to improve the quality of life of PDAC patients as current chemotherapy regimens all have major toxicities. Furthermore, we demonstrated that the survival time of mice treated with 5-FU-miR-15a alone or in combination with gemcitabine was significantly longer than control. While gemcitabine alone was more effective for extending survival than 5-FU-miR-15a alone, the dose of gemcitabine was 15 fold higher than 5-FU-miR-15a. The effects of 5-FU-miR-15a on survival demonstrate its therapeutic potential. Future studies are needed to optimize the therapeutic dose and treatment frequency to improve the impact on survival. These findings are consistent with our *in vitro* data and demonstrate the advantage of a multi-targeted therapeutic that inhibits the expression of several important oncogenes. As delivery is a major challenge for miRNA therapeutics, for this experiment we used *in vivo-*JetPEI as the delivery vehicle. *In vivo-*JetPEI has been demonstrated to be a highly effective and non-toxic agent for siRNA and miRNA drug development ^48^. Taken together, our results show potential for 5-FU-miR-15a mimic as a novel therapeutic agent alone or in combination with the current treatment regimen in treating PDAC. This is the first study to demonstrate the effectiveness of a 5-FU modified miRNA as a potential novel therapy for treating metastatic PDAC via systemic delivery. While the 5 year survival for all pancreatic cancer patients is very low, for patients with metastatic disease it is only 3% demonstrating the important need for novel therapeutics that are effective for treating metastatic disease ^1^.

In summary, our study clearly defines the molecular and functional significance of miR-15a as a tumor suppressor in PDAC by suppressing the expression of several key targets (Wee1, Chk1, BMI-1, and Yap-1). miR-15a abrogates the G2 cell cycle check point control by suppressing the expression of Wee1 and Chk1. miR-15a also has potential to be a prognostic biomarker in future clinical management of PDAC. We demonstrated that 5-FU integrated miR-15a mimic has potent therapeutic power for inhibiting PDAC lung metastatic tumor growth without any observed toxic side effects. The strategy of incorporating 5-FU into miR-15a to create a new more effective therapeutic entity will be a platform technology for creating miRNA based therapeutics for treating PDAC as well as other tumor types.

## Materials and Methods

### Cell culture

The human pancreatic cancer cell lines (AsPC-1, PANC-1 and Hs 766T) were purchased from the American Type Culture Collection (ATCC, Manassas, VA, USA), and maintained in RPMI (AsPC-1) or DMEM (PANC-1 and Hs 766T), (Thermo Fisher, Waltham MA) medium supplemented with 10% fetal bovine serum (Thermo Fisher).

### Tissue samples

Thirty fresh-frozen pancreatic cancer specimens and ten fresh-frozen normal pancreas tissues were obtained from the Institute of Hepatopancreatobiliary Surgery, Southwest Hospital, Third Military Medical University (Army Medical University). Ten normal pancreatic samples were obtained from organ donors. One hundred and eleven formalin-fixed paraffin-embedded (FFPE) tissues (including 101 pancreatic cancer tissues and 10 normal pancreas tissues) were acquired from the archival collections of Southwest Hospital, Third Military Medical University and the First Affiliated Hospital of Soochow University. None of these patients had received preoperative adjuvant chemotherapy or radiotherapy. The use of human tissues was approved by the ethics committee of Southwest Hospital, Third Military Medical University (Army Medical University), Chongqing, China.

### TCGA survival analysis

We obtained the clinical and miRNA expression data for TCGA pancreatic ductal adenocarcinoma from the UCSC cancer genome browser^49^. There are 176 available cases with both survival and miR-15a/Cdc2 expression data in TCGA database. Based on dichotomizing miR-15a expression profile, we divided cases into high and low expression groups. The cut-off is 50%, that is, expression greater than the 50 percentile of expression of the patients was denoted as “high”; otherwise it was denoted as “low”. Subsequently, we performed log-rank tests between two groups and obtained corresponding Kaplan–Meier curves.

### Transfection

Twenty four hours before transfection, 2×10^5^cells per well were plated in a six-well plate and transfected with 50 nM of either miR-15a, non-specific control miRNA (Thermo Fisher) or modified 5-FU-miR-15a (Dharmacon Lafayette, CO) using Oligofectamine (Thermo Fisher) following the manufacturer’s protocols.

### Cell proliferation

Twenty-four hours after transfection, 2000 cells per well were replated in a 96-well plate. Cell viability was assessed on days 1, 3 and 6 post transfection using WST-1 dye (Roche, Basel, Switzerland) as previously described ^17^.

### Cell cycle analysis

Twenty-four hours after transfection, cells were collected and resuspended in modified Krishan buffer containing 0.02 mg/ml RNase H (Thermo Fisher) and 0.05 mg/ml Propidium Iodide (Sigma Aldrich, St. Louis, MO). Stained cells were detected by flow cytometry. The data was analyzed using ModFit software (BD Bioscience, Sparks, MD).

### Western blotting analysis

Equal amounts of protein (15 μg) were separated on 10%-12% sodium dodecyl sulfate polyacrylamide gels by the method of Laemmli ^50^. Proteins were probed with anti-Yap-1 monoclonal antibody (1:10000) (Cell Signaling Technologies, Danvers, MA), anti-BMI-1 (1:10000) (Cell Signaling Technologies), anti-Wee1 (1:500) (Cell Signaling Technologies), anti-Chk1 (1:500) (Cell Signaling Technologies), anti-Cdc2 (1:500) (Cell Signaling Technologies), anti-p27 (1:500) (Santa Cruz Biotech Inc., Santa Cruz, CA), anti-Cyclin A (1:500) (Santa Cruz Biotech Inc.), anti-GAPDH (1:200000) (Santa Cruz Biotech Inc.). Horseradish peroxidase conjugated antibodies against mouse or rabbit (1:5000, Santa Cruz Biotech Inc.) were used as the secondary antibodies. Protein bands were visualized using a LI-COR Odyssey FC Imaging system and Super Signal West Pico Chemiluminescent Substrate (Thermo Fisher). Western blots were quantified using Image Studio Software. All Western blots were repeated 3 times for quantification.

### Luciferase reporter assay

Luciferase reporter plasmids containing the Wee1 mRNA3’-UTR (HmiT054534-MT06) or the Chk1 mRNA3’-UTR (HmiT061994-MT06) were purchased from GeneCopoeia. Twenty four hours before transfection, 1.0×10^4^cells were plated in 96-well plate. 10 nM of miR-15a/5-FU-miR-15a or control miRNA was transfected into these cells together with 100 ng of pEZX-MT06-Report-Wee1 or pEZX-MT06-Report-Chk1 by DharmaFect Duo (Dharmacon) following the manufacturer’s protocol. The luciferase assay was performed 24 hours after transfection using a dual-luciferase reporter assay system (Promega, Madison, WI). For each sample, firefly luciferase activity was normalized to Renilla luciferase activity and to control miRNA transfected cells.

### RNA isolation and real-time qRT-PCR

Total RNA from the transfected pancreatic cancer cell lines as well as pancreatic cancer specimens was extracted using TRIzol reagent. For miR-15a and miR-16 detection in clinical samples, cDNA synthesis was performed using a PrimeScript RT reagent kit (TaKaRa, Kusatsu, Shiga Prefecture, Japan), and qRT–PCR was performed with SYBR Premix Ex Taq II (TaKaRa). The expression levels were normalized to the endogenous small nuclear RNA U6 control. For primary, precursor, mature miR-15a and miR-16 detection in pancreatic cancer cell lines and pancreatic cancer tissues, primary, precursor and mature miR-15a/miR-16 specific primers were purchased from Thermo Fisher. cDNA synthesis was performed by the High Capacity cDNA Synthesis Kit (Thermo Fisher) with miRNA-specific primers. Real-time qRT-PCR was carried out using miRNA-specific primers by TaqMan Gene Expression Assay (Thermo Fischer). Expression of precursor and mature was normalized to that of the primary transcript for analysis. The separation of nuclear and cytoplasmic fractions was performed using PARIS^™^ Kit (Thermo Fisher).

### Gemcitabine and cytotoxicity assay

Twenty four hours after transfection, Hs 766T cells were replated in a 96-well plate at 2000 cells per well in triplicate, in 100μl of medium supplemented with 10% Dialyzed FBS (Thermo Fisher). After 24 hours, fresh media containing gemcitabine alone (ranging from 0 to 20 μM), 5-FU-miR-15a alone (ranging from 0 to 200 nM) or 5-FU-miR-15a and gemcitabine in combination (at a constant ratio 1:100, with increasing concentrations of both compounds) was added, and cells were cultured for 72 hours. Cell viability was measured using the WST-1 assay, and concentration dependent curves were generated based on the cell viability. The Combination Index (CI) as well as IC^50^ for each alone or in combination was calculated using CompuSyn software (www.combosyn.com).

### Immunohistochemistry (IHC)

IHC analysis was performed using a standard streptavidin–biotin-peroxidase complex method. Briefly, the slides were incubated with Cdc2 primary antibodies (Cell Signaling Technologies) overnight, followed by incubation with secondary antibodies and further incubation with the streptavidin–biotin complex (Maixin, Fuzhou, China). Cdc2-positive cells were defined as those with brown staining in the nucleus and cytoplasm, and the expression was assessed based on the percentage of positive tumor cells out of 1000 tumor cells. The quantification was performed by using a composite score obtained by multiplying the values of staining intensities (0, no staining; 1, weak staining; 2, moderate staining; 3, strong staining) and the percentage of positive cells (0, 0%; 1, <10%; 2, 10–50%; 3, >50%). For statistical analysis, the tumor sample cohort was grouped into those with low expression (≤4) and those with high expression (≥ 6).

### Mouse pancreatic cancer metastasis model

All animal procedures were approved by the Stony Brook University Institutional Animal Care and Use Committee (IACUC). NOD/SCID mice were purchased from Jackson Lab. For evaluating the impact of 5-FU-miR-15a in a pancreatic cancer metastasis model, mouse tumors were established via tail vein injection of 2×10^6^ Hs 766T luciferase expressing cells into 24, 8 week old NOD/SCID mice. Mice were divided into four groups (control, gemcitabine alone, 5-FU-miR-15a alone and 5-FU-miR-15a plus gemcitabine) six mice per group. Three days after injection, mice were treated via tail vein injection with 80 μg of control miRNA or 5-FdU-miR-15a packaged using *in vivo*-jetPEI (Polyplus Transfection, Illkirch, France) or with 50 mg/kg of gemcitabine (Sigma Aldrich) via intraperitoneal injection (IP injection). In the first stage, mice were treated with 5-FU-miR-15a every other day for 2 weeks (8 times) and gemcitabine every three days for 2 weeks (4 times). Following treatment, mice were screened using IVIS Spectrum In vivo Imaging System (IVIS) (PerkinElmer, Waltham, MA) for 2 weeks. Subsequently, treatment was restarted for one week including 5-FU-miR-15a (4 times) and gemcitabine (2 times), and then screened using IVIS again. Mice were monitored daily to determine the survival time.

### Statistical analysis

All experiments were repeated at least three times. All statistical analyses were performed with GraphPad Prism Software. The statistical significance between two groups was determined using Student’s t-test. For comparison of more than two groups, one-way ANOVA followed by a Bonferroni-Dunn test was used. Data are expressed as mean ± standard error of the mean (SEM). The statistical significance is either described in figure legends, or indicated with asterisks (*p<0.05;**p<0.01; ***p<0.001).

## Acknowledgments

This study was supported by grants from National Key R&D Program of China (No. 2017YFC1308600), the National Institute of Health/National Cancer Institute R01CA15501904 (J Ju), National Natural Science Foundation of China (No. 81502550) and Southwest Hospital Research Project (No. SWH2016JCYB-45). A.F. and J.J. have filed a patent for 5-FU-miR-15a, all other authors declare that they have no competing interests.

## Author Contributions

S.G. conceived and performed experiments, analyzed data and wrote the manuscript. A.F. conceived of and directed experiments, performed animal experiments and revised the manuscript. W.H. performed real-time PCR for miR-15a/miR-16 in clinical samples. Y.W. and J.Y. collected and summarized the clinical information and follow-up documents for PDAC patients. X.W. and Y.Z. performed IHC for CDC2 in clinical samples. G.H. helped to perform IVIS Spectrum for vivo experiment. H.Z. and J.J. conceived and supervised the project, acquired funding, and reviewed and edited the manuscript. All authors read and approved the final manuscript.

## Supplemental Figures

**Figure S1.**
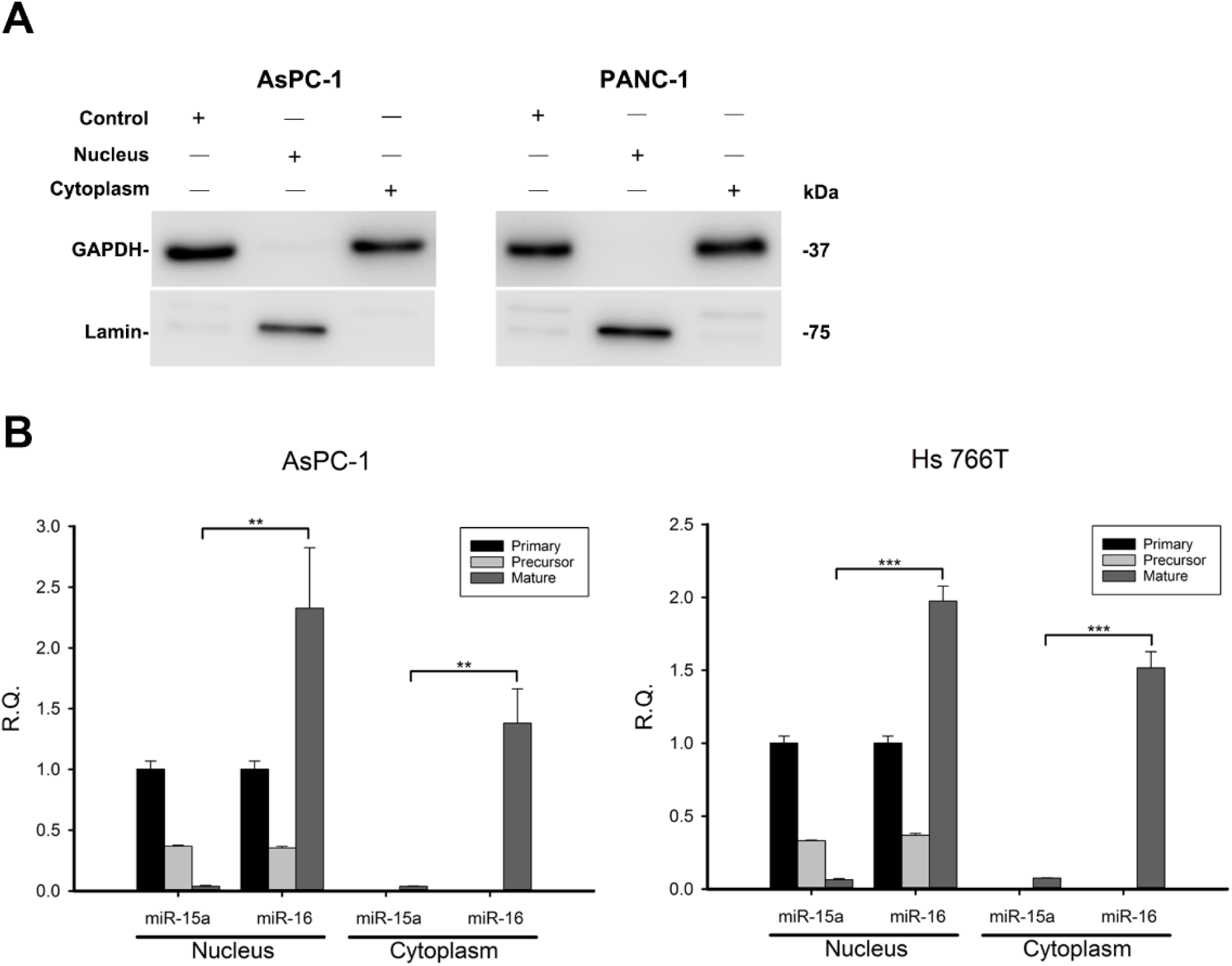
Processing of miR-15a and miR-16 in PDAC. **(A)** Western blot showing nuclear and cytoplasmic separation using the PARIS Kit **(B)** In AsPC-1 and Hs 766T cells expression of pre-miR-15a and pre-miR-16 are similar in both the nucleus and the cytoplasm. Expression of mature miR-16 is significantly higher than miR-15a in both the nucleus and the cytoplasm. (** p<0.01, *** p<0.001)

**Figure S2.**
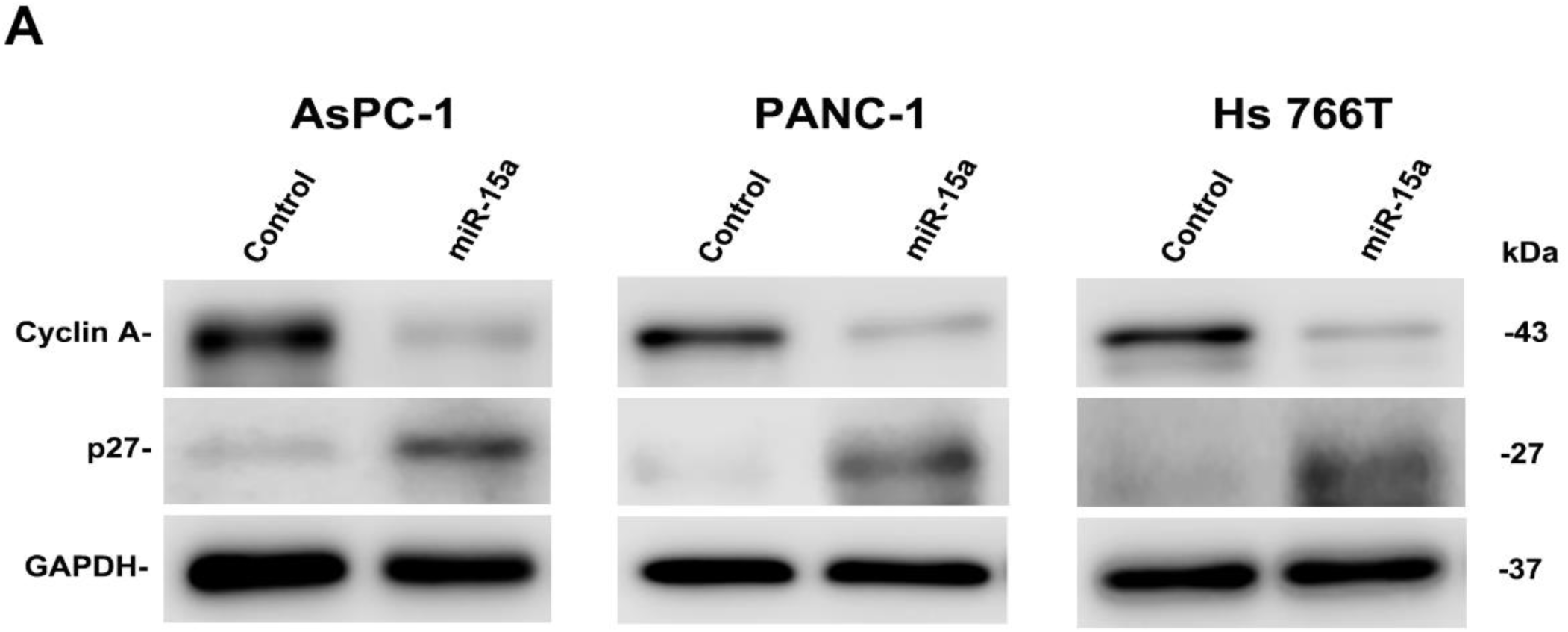
miR-15a induces cell cycle arrest. (**A**) miR-15a regulates the expressions of downstream targets correlated with G1 arrest (p27, Cyclin A)

**Figure S3.**
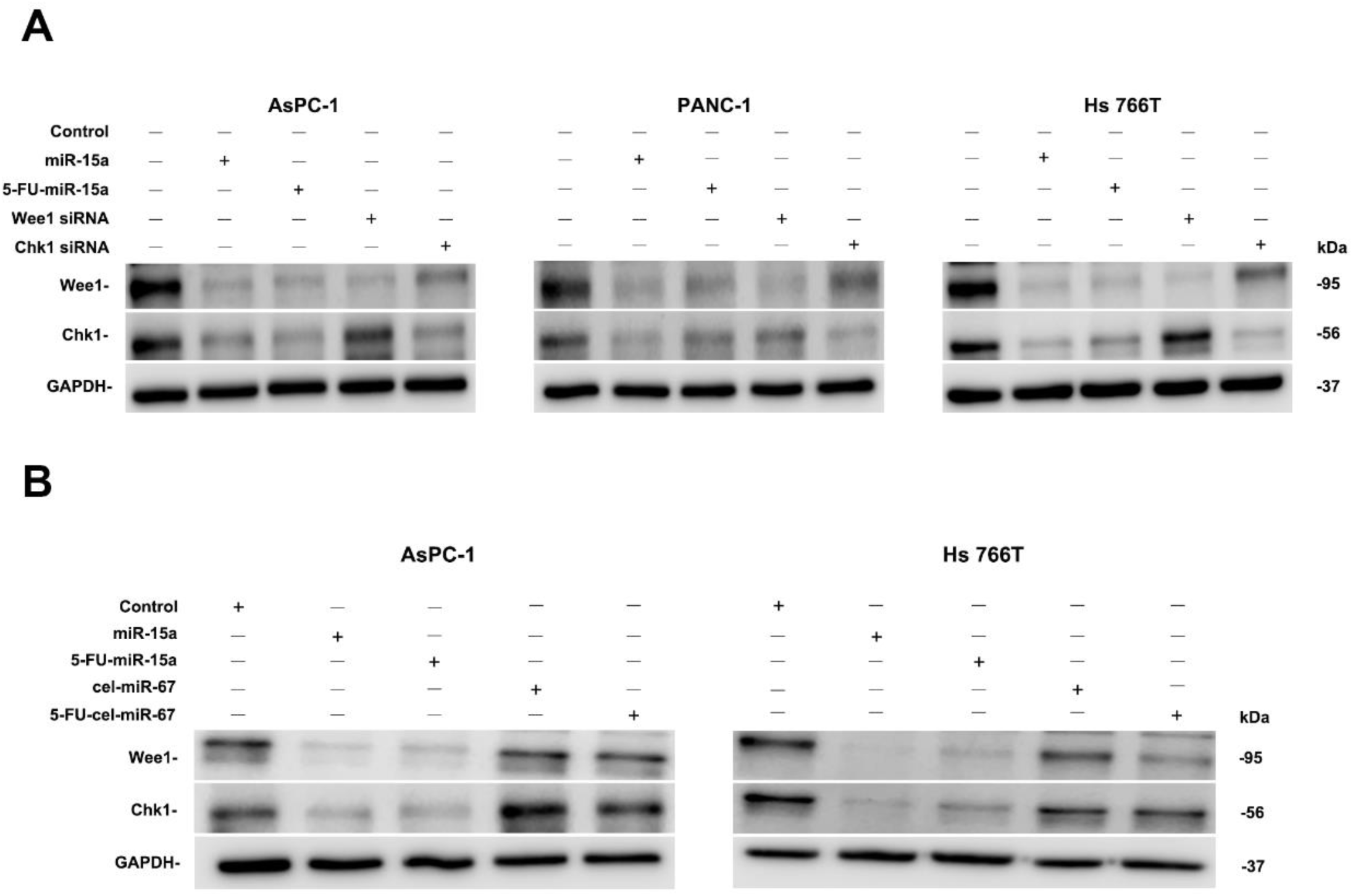
miR-15a inhibits several important targets in PDAC. **(A)** Wee1/Chk1expression levels of cells transfected with Wee1 siRNA or Chk1 siRNA are consistent with that of cells transfected with 5-FU-miR-15a. **(B)** 5-FU-miR-15a significantly decreased the expressions of Wee1 and Chk1 compared to 5-FU-cel-miR-67.

